# Humans exploit robust locomotion by improving the stability of control signals

**DOI:** 10.1101/625855

**Authors:** Alessandro Santuz, Leon Brüll, Antonis Ekizos, Arno Schroll, Nils Eckardt, Armin Kibele, Michael Schwenk, Adamantios Arampatzis

## Abstract

Is the control of movement less stable when we walk or run in challenging settings? One might intuitively answer affirmatively, given that adding constraints to locomotion (e.g. rough terrain, age-related impairments, etc.) imply less stable movements. We investigated how young and old humans synergistically activate muscles during locomotion, when different perturbation levels are introduced. Of these control signals, called muscle synergies, we then analyzed the stability over time. Surprisingly, we found that perturbations and older age force the central nervous system to produce more stable signals. These outcomes show that robust locomotion in challenging settings is achieved by increasing the stability of control signals, whereas easier tasks allow for more unstable control.

## Main Text

The central nervous system (CNS), as fundamental nonlinear component of the majority of animals, is intrinsically chaotic (*1*) and noisy (*2*). A chaotic system must be indecomposable into subsystems, with “an element of regularity”, though unpredictable (*3*). In mathematical terms, it must be topologically mixing (or transitive) and have dense periodic orbits, conditions which imply that it is sensitive to initial conditions (*4*). In other words, small variations in the initial conditions might generate big variations in the evolution of the system, even though these deviations are not random but fully determined by the initial conditions themselves (*5*). Noise affects neural control signals by adding random or irregular disturbances in a signal-dependent manner: if the magnitude of the control signal increases, noise levels increase as well (*6*). To organize reliable movements, the CNS must robustly handle chaotic and noisy elements (*1*, *2*). Defining robustness as the ability to cope with perturbations (*7*), it follows that biological systems can manage to maintain function despite disturbances only through robust control (*8*–*10*). Assessing the stability, which is the sensitivity to infinitesimally small perturbations (*5*), of control signals could give us an idea of the strategies adopted by the CNS to robustly cope with disruptions, whether they are internal (e.g. aging or disease) or external (e.g. environmental, such as changes in the morphology of terrain).

With this study, we propose an innovative approach to describe the dynamic stability (*5*, *11*, *12*) of human motor control applied to locomotion. We started from a crucial question: how is the stability of control signals associated to robust motor output? The answer might give important insight into the neural mechanisms necessary for the robust control of vertebrate locomotion. Despite chaos and noise which add to the overall non-linearity, the output of the CNS can be reasonably described and modelled by means of linear approximations (*13*). The overwhelming amount of degrees of freedom available to vertebrates for accomplishing any kind of movement is defined by the vast number of muscles and joints. Yet, the CNS manages to overcome complexity, possibly through the orchestrated activation of functionally-related muscle groups, rather than through individual entities (*14*, *15*). These common activation patterns, called muscle synergies, are thought to be used by the CNS for simplifying the motor control problem by reducing its dimensionality (*13*). Usually extracted from electromyographic (EMG) data by means of linear machine learning approaches such as the non-linear matrix factorization (NMF), muscle synergies have been increasingly employed in the past two decades for providing indirect evidence of a simplified, modular control of movement in humans and other vertebrates (*13*, *16*–*18*).

We quantified the dynamic stability of motor primitives (i.e. the temporal components of muscle synergies) in different locomotor tasks and settings by means of the maximum finite-time Lyapunov exponents (MLE), a metric used to describe the rate of separation of infinitesimally close trajectories (*19*). Namely, we considered what happens in the space of muscle synergies when humans switch from walking to running, from overground to treadmill locomotion, from unperturbed to perturbed locomotion and in aging. We chose these conditions to compare different kinds of constraints on locomotion: running allows less time for organizing coordinated movements than walking (*20*); externally-perturbed locomotion is more constrained than unperturbed due to the increased mechanical and physiological limits imposed by the higher environmental complexity (*7*, *21*); aging is a source of internal perturbation which leads to muscle weakness and loss of fine neural control at various levels (e.g. CNS, proprioceptive, etc.) (*22*, *23*). Recently, we and others proposed that the width of motor primitives increases to ensure robust control in the presence of internal and external perturbations (*7*, *24*–*27*). We observed this neural strategy in wild-type mice (*27*) and in humans (*7*) undergoing external perturbations, but not in genetically modified mice that lacked feedback from proprioceptors (*27*). Due to these observations, we concluded that intact systems use wider (i.e. of longer duration) control signals to create an overlap between chronologically-adjacent synergies to regulate motor function (*7*). Our conclusions, however, did not include any information about the stability of the control signals, eventually limiting the understanding of the adaptive processes needed to cope with perturbations.

With the present analysis of human data, we discovered that more stable motor primitives are associated with more constrained conditions, whereas easier, less constrained tasks or younger age allow for more unstable control. Our findings provide new important insights on how our CNS might control repetitive, highly-stereotyped movements like locomotion, in the presence and absence of perturbations. Moreover, these results add interesting aspects to the definition of robust motor control that we and others (*7*, *28*) gave in the past, integrating the concept of stability with the width of control signals.

## Results

Muscle synergies for human locomotion have been extensively discussed and reported in the past (*7*, *29*–*32*). Fig. 1 shows a typical output for walking, where four fundamental synergies describe as many phases of the gait cycle. In human locomotion, the first synergy functionally refers to the body weight acceptance, with a major involvement of knee extensors and glutei. The second synergy describes the propulsion phase, to which the plantarflexors mainly contribute. The third synergy identifies the early swing, showing the involvement of foot dorsiflexors. The fourth and last synergy reflects the late swing and the landing preparation, highlighting the relevant influence of knee flexors and foot dorsiflexors.

**Fig. 1.**
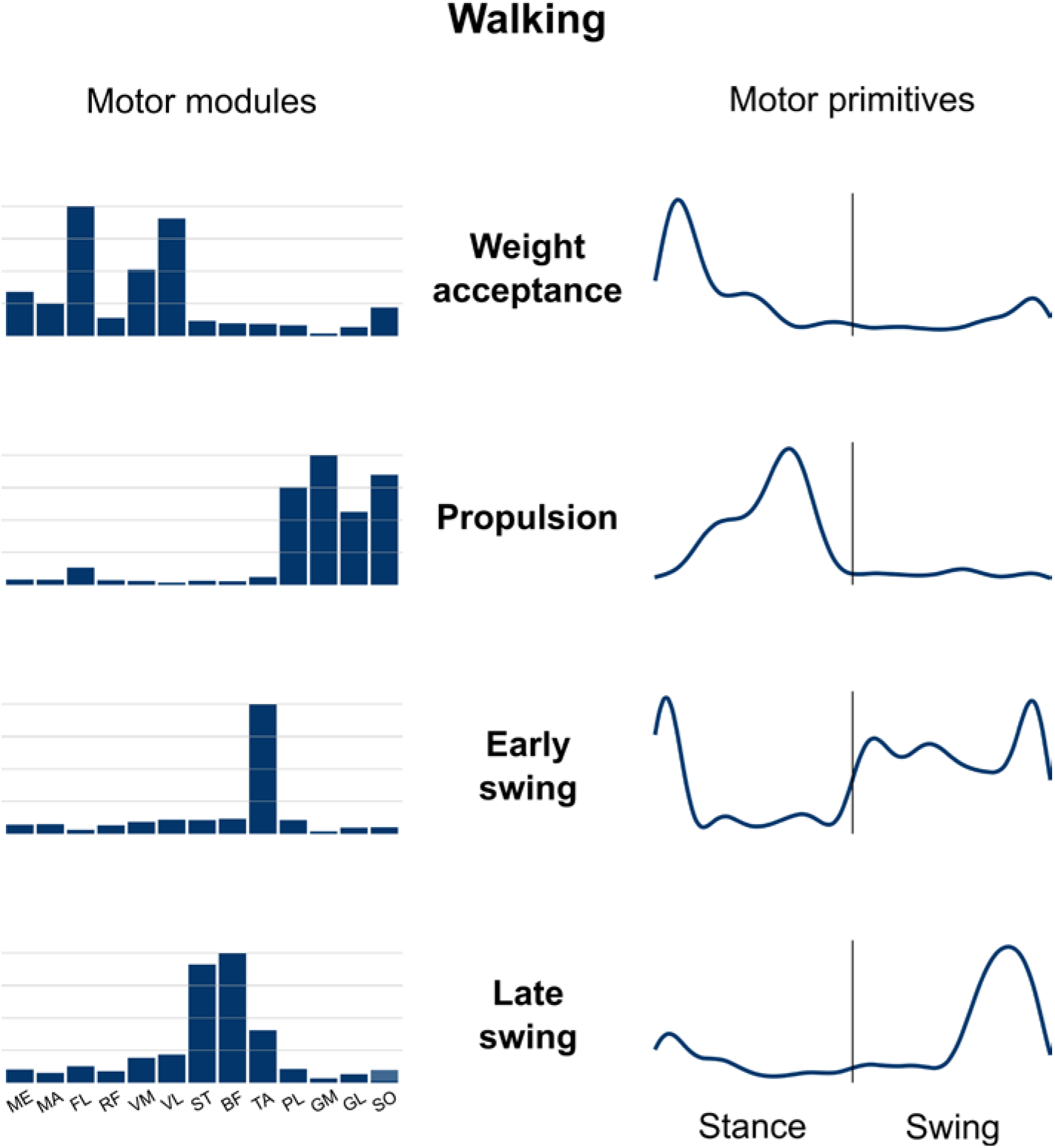
Muscle synergies for human walking. Exemplary motor modules and motor primitives of the four fundamental synergies for human walking (average of even-surface trials of the experimental setup E2). The motor modules are presented on a normalized y-axis base. For the motor primitives, the x-axis full scale represents the averaged gait cycle (with stance and swing normalized to the same amount of points and divided by a vertical line) and the y-axis the normalized amplitude. Muscle abbreviations: MA=gluteus maximus, ME=gluteus medius, FL=tensor fasciæ latæ, RF=rectus femoris, VM=vastus medialis, VL=vastus lateralis, ST=semitendinosus, BF=biceps femoris, TA=tibialis anterior, PL=peroneus longus, GM=gastrocnemius medialis, GL=gastrocnemius lateralis, SO=soleus.

Details about the participants are reported in the supplementary materials of this paper. Briefly, the first group (G1) of young participants was assigned to the first experimental protocol (E1) consisting of walking and running overground and on a treadmill. The second group (G2) was assigned to the second experimental protocol (E2) consisting of walking and running on one standard and one uneven-surface (Fig. S1, Movie S1) treadmill. The last two groups (G3, young and G4, old) where assigned to the third and last protocol (E3) which consisted of walking on a treadmill (Movie S1) that could provide mediolateral and anteroposterior perturbations. The minimum number of synergies which best accounted for the EMG data variance (i.e. the factorization rank) of E1 was 4.6 ± 0.5 (G1, overground walking), 4.6 ± 0.6 (G1, treadmill walking), 4.2 ± 0.7 (G1, overground running), and 4.6 ± 0.7 (treadmill running), with no significant main effects of locomotion type or condition. In E2, the values were 4.7 ± 0.7 (G2, even-surface walking), 5.1 ± 0.6 (G2, uneven-surface walking), 4.2 ± 0.6 (G2, even-surface running), and 4.6 ± 0.6 (G2, uneven-surface running), with running showing significantly less synergies than walking (p = 0.005) and no statistically significant effect of surface. Finally, in E3 the factorization ranks were 4.5 ± 0.6 (G3, unperturbed walking, young), 4.9 ± 0.5 (G3, perturbed walking, young), 4.5 ± 0.6 (G4, unperturbed walking, old), and 4.6 ± 0.5 (G4, perturbed walking, old), with perturbed walking in the young showing significantly more synergies (p < 0.001).

In Table 1 we report, synergy-by-synergy and for the three experimental setups, the factors which contributed to widen the motor primitives. Detailed boxplots are available in Fig. S2. Our findings show that a widening of the motor primitives can be observed in a) running compared to walking, b) perturbed compared to unperturbed locomotion and c) old compared to young participants (Table 1).

**Table 1.**
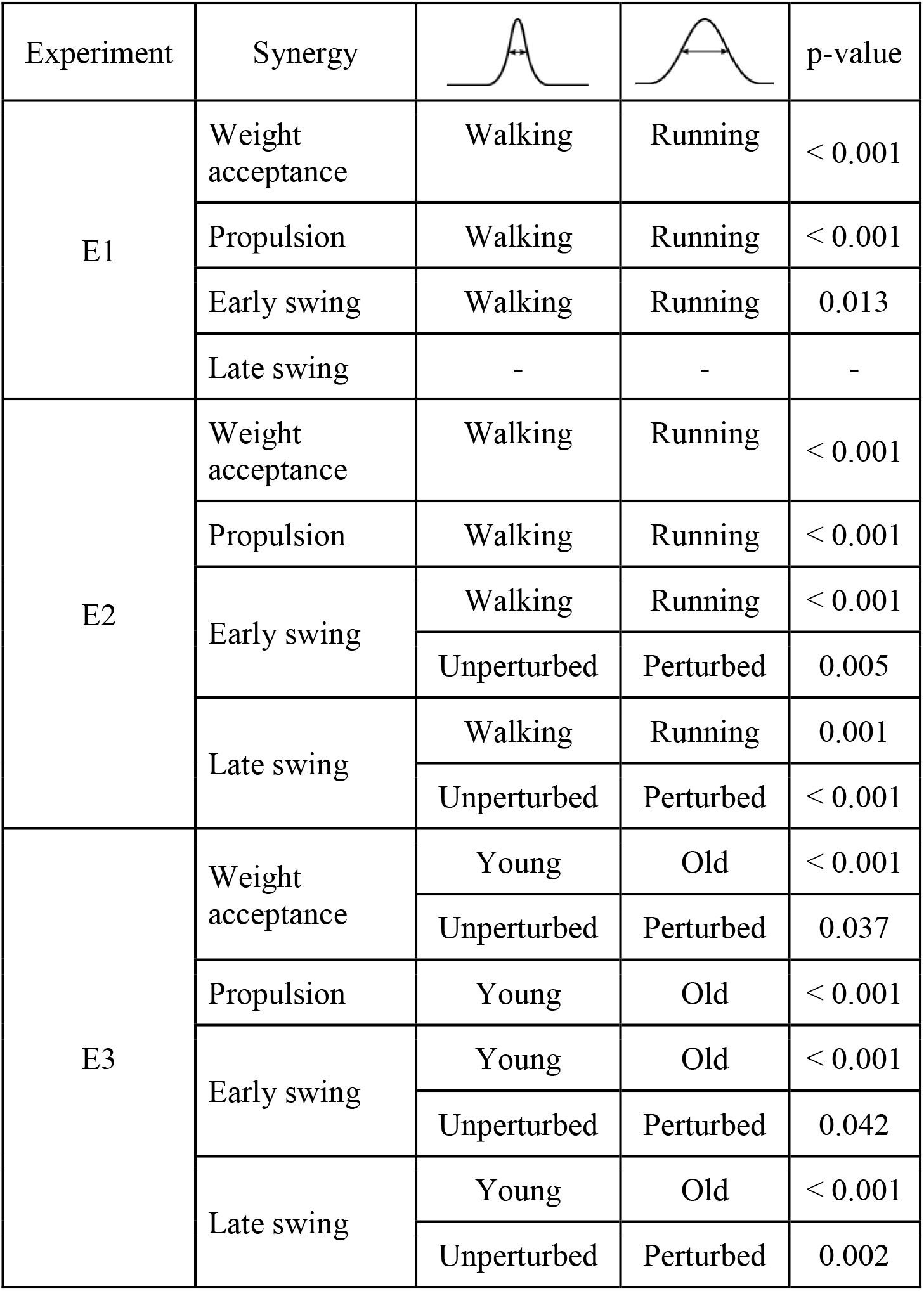
Widening of motor primitives. Summary of the conditions that had an effect on the full width at half maximum of the motor primitives extracted from the data of the three experimental setups (E1 = walking and running, overground and treadmill; E2 = walking and running, even- and uneven-surface; E3 = unperturbed and perturbed walking, young and old). Motor primitives are the temporal coefficients of the four fundamental synergies for locomotion. Detailed boxplots are available in Fig. S2.

| Experiment | Synergy |  |  | p-value |
| --- | --- | --- | --- | --- |
| E1 | Weight acceptance | Walking | Running | < 0.001 |
|  | Propulsion | Walking | Running | < 0.001 |
|  | Early swing | Walking | Running | 0.013 |
|  | Late swing | - | - | - |
| E2 | Weight acceptance | Walking | Running | < 0.001 |
|  | Propulsion | Walking | Running | < 0.001 |
|  | Early swing | Walking | Running | < 0.001 |
|  |  | Unperturbed | Perturbed | 0.005 |
|  | Late swing | Walking | Running | 0.001 |
|  |  | Unperturbed | Perturbed | < 0.001 |
| E3 | Weight acceptance | Young | Old | < 0.001 |
|  |  | Unperturbed | Perturbed | 0.037 |
|  | Propulsion | Young | Old | < 0.001 |
|  | Early swing | Young | Old | < 0.001 |
|  |  | Unperturbed | Perturbed | 0.042 |
|  | Late swing | Young | Old | < 0.001 |
|  |  | Unperturbed | Perturbed | 0.002 |

We analyzed motor primitives in their own space, the dimension of which was equal to the trial-specific number of synergies. Two representative trials factorized into three synergies each are plot in 3-dimensional graphs in Fig. 2. We used the 3-dimensional example since a space in more than three dimensions would be difficult to represent graphically. We found that motor primitives are more stable (i.e. show lower MLE) in a) running compared to walking, b) perturbed compared to unperturbed locomotion and c) in old compared to young participants (Fig. 3 and S3). Motor primitives did not show any difference in the MLE when comparing overground with treadmill locomotion.

**Fig. 2.**
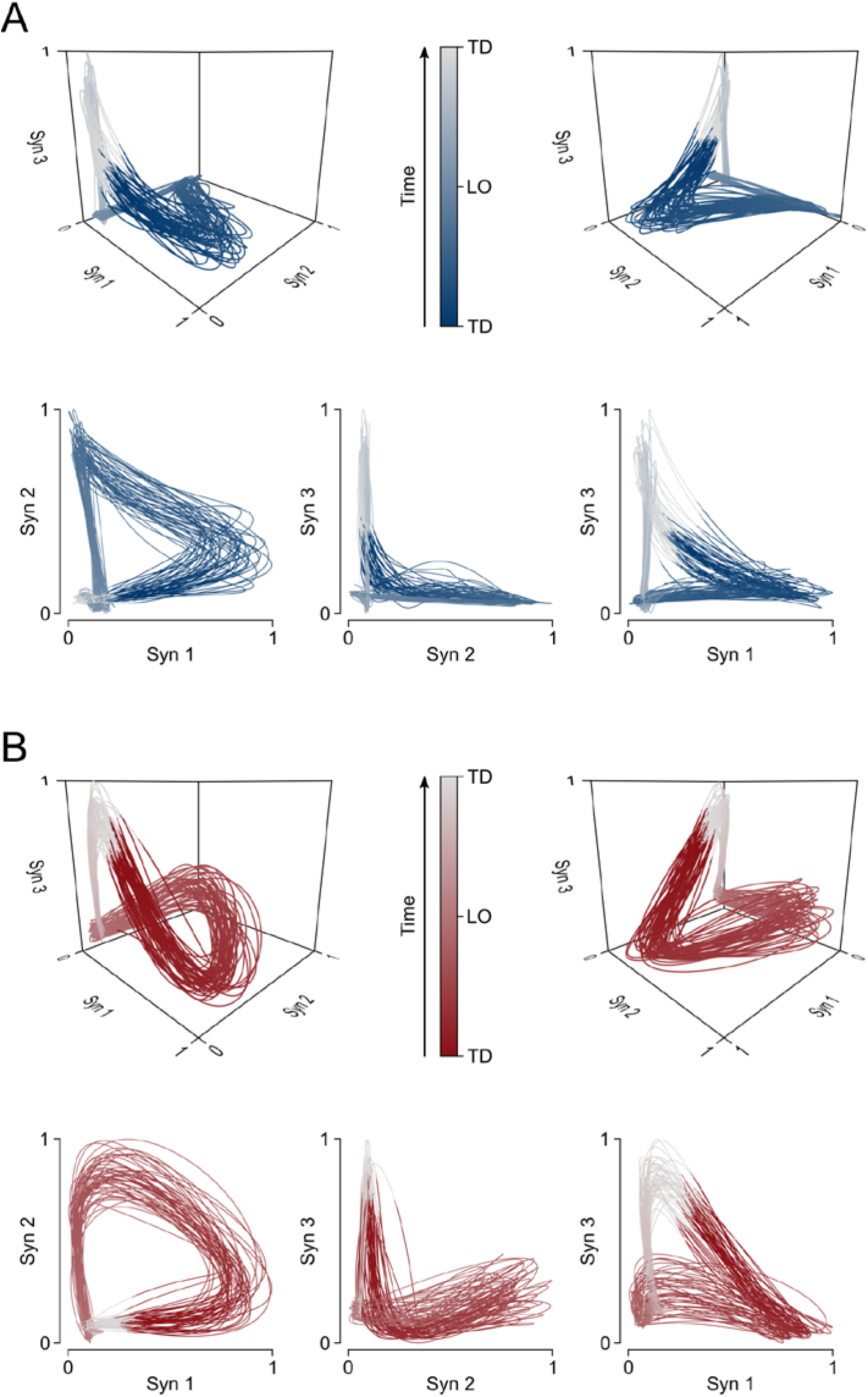
Motor primitive trajectories in their own space. Representative data showing the filtered trajectories of motor primitives when the number of synergies (Syn) is equal to three. Panel A (blue curves) refers to a walking trial, unperturbed, young participant. Panel B (red curves) refers to a running trial, unperturbed, young participant. Trajectories are color-coded from touchdown (TD, dark blue or red), to lift-off (LO, light blue or red), to the next TD (white). The amplitude of motor primitives is normalized to the maximum value of each trial for better visualization.

**Fig. 3.**
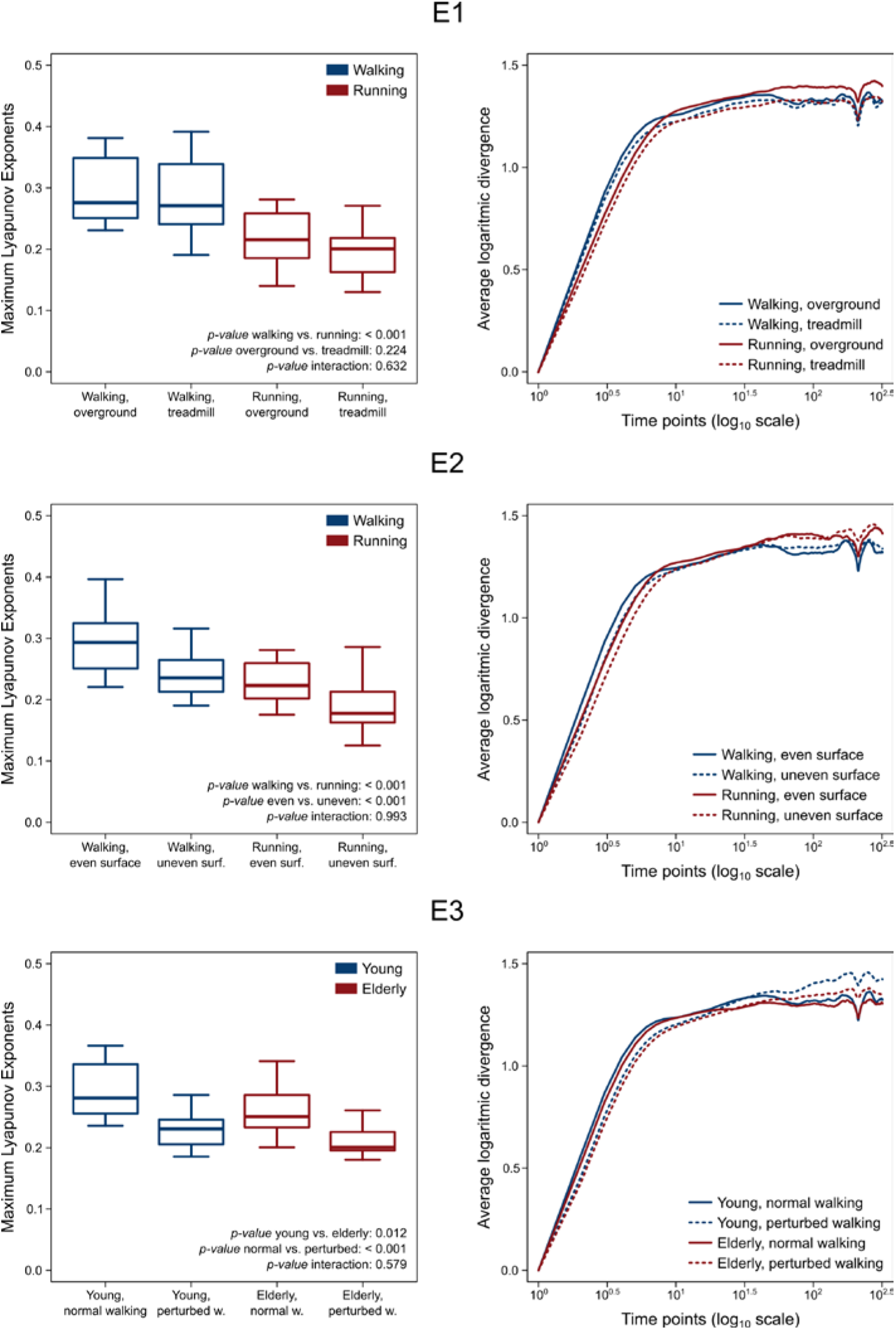
Maximum Lyapunov Exponents of motor primitives. Boxplots and curves describing the maximum Lyapunov exponents (MLE) and the average logarithmic divergence curves for the three experimental setups (E1 = walking and running, overground and treadmill; E2 = walking and running, even- and uneven-surface; E3 = unperturbed and perturbed walking, young and old). The minimum value was subtracted from each curve for improving the visualization. The actual vertical intercept was negative and different for all curves (Fig. S3). Lower MLE imply more stable motor primitives.

## Discussion

MLE contain information about chaoticity and noisiness (*11*, *12*), both of which are intrinsic properties of neural systems (*1*, *2*). Chaos is thought to be required or “almost unavoidable” for exploring the available opportunities of action when the system is challenged and, consequently, to maintain robustness, which is the ability to cope with perturbations (*1*, *7*). The presence of internal and external noise, together with the resulting constraints, shaped the evolution of neural systems and might be beneficial for information processing (*2*). Yet, whether different levels of robustness imply different stability of neural control is a question that does not find intuitive answers. Historically, MLE have been used to give information about the behavior of chaotic dynamical systems (*7*, *11*, *20*, *33*, *34*). In this study, we described the stability of modular motor control in humans by calculating the MLE of motor primitives (i.e. the time-dependent coefficients of muscle synergies) during locomotion (walking and running) overground and on a treadmill, with or without external perturbations and in aging. Our results show higher stability (i.e. lower MLE) and longer basic activation patterns (i.e. higher FWHM) associated with aging, external perturbations, and the switch from walking to running. We proposed an innovative and simple approach to describe the behavior of neural system modularity, with an eye on increasing the reproducibility of results. Typically, MLE are calculated from data expanded in the state space, which is a set of all the possible states of a system at any given time (*1*, *5*, *35*). The main assumption underlying our method is that the analysis must be conducted in the muscle synergies space, with its own dimension which is equal to the factorization rank (i.e. the minimum number of synergies necessary to sufficiently reconstruct the original EMG signals). By doing so, we did not model the whole system dynamics, but focused on the modular behavior of the CNS. In this assumption lies also the high reproducibility of our approach, since this simplification of the calculations avoids a well-known weakness (*12*) of the classical approach: the choice of time delay and state space dimension for delay embedding.

In the past, we used the FWHM of motor primitives as a measure of robustness (*7*). Our conclusion was that wider (i.e. timewise longer active) primitives indicate more robust control (*7*). We reasoned that the overlap of chronologically-adjacent synergies increased the fuzziness (*9*, *36*) of temporal boundaries allowing for easier shifts between one synergy (or gait phase) to the other (*7*), a conclusion that fits the optimal feedback control theory (*37*, *38*). For the CNS, this solution must come at a cost: the reduction of accuracy or, as others called it, optimality (*9*) or efficiency (*39*). For instance, it has been recently found that human neurons allow less vocabulary overlap than monkey’s, showing a tradeoff between accuracy (complex human feature) and robustness (basic, typical of non-human primates) across species (*39*). In this study we confirmed a widening of motor primitives in those conditions that were more constrained than their equivalent baseline and in aging. Specifically, we considered running as a more constrained locomotion type than walking (*20*), treadmill-as more constrained than overground-locomotion (*11*) and perturbed-as more constrained than unperturbed-locomotion (*7*). We found an effect of motor constraints and aging on the widening of motor primitives.

However, we discovered that aging and the more constrained locomotion conditions imply not only wider primitives, but a different stability of neural control as well. We calculated lower MLE (i.e. higher stability) in old compared to young, in running compared to walking and in perturbed compared to unperturbed locomotion. These outcomes indicate that robustness is not only achieved by allowing motor primitives to be wider and fuzzier, but by making them more stable as well. Interestingly, we recently found that the classical calculation of MLE from kinematic data (e.g. by considering the trajectories of specific body landmarks recorded via motion capture) shows decreased stability in the presence of constraints in both humans (*7*, *20*, *33*) and mice (*27*). Our interpretation of this apparent discordance lies in the results of the present study. The MLE calculation by means of state space reconstruction acts as a representation of the human locomotor system as a whole (*11*). Thus, increased MLE mean higher sensitivity of the entire dynamical system to infinitesimal perturbations (*11*). The analysis we propose, though, does not include any sort of state space embedding, and aims to the description of the CNS as a subsystem for the control of the main system’s motion. This rationale tells us that the two descriptions are intrinsically different, possibly because they describe different portions of the “human being” as a dynamical system. From this perspective, it is not surprising that two different approaches give opposite results. In fact, the higher stability of motor primitives might describe a strategy employed by the CNS to maintain acceptable levels of functionality when constraints are added globally.

In conclusion, our analysis reveals a new view on neural stability: fuzzier, more robust muscle activation patterns are generated by the CNS in the presence of constraints to cope with perturbations (*7*). The stability of neural control increases when constraints are added to movement, ensuring robust locomotion across a variety of challenges.

## Materials and Methods

This study was reviewed and approved by the Ethics Committees of the Humboldt-Universität zu Berlin, Kassel University and Heidelberg University. All the participants gave written informed consent for the experimental procedure, in accordance with the Declaration of Helsinki.

### Experimental protocols

For the three experimental protocols we recruited 86 healthy volunteers and divided them into four groups. The first group of 30 (henceforth G1, 15 males and 15 females, height 173 ± 10 cm, body mass 68 ± 12 kg, age 27 ± 5 years, means ± standard deviation) was assigned to the first experimental protocol (E1). The second group of 18 (G2, 11 males and 7 females, height 176 ± 7 cm, body mass 71 ± 13 kg, age 24 ± 3 years) was assigned to the second experimental protocol (E2). The last two groups where assigned to the third and last protocol (E3): one group of young (G3, 7 males and 12 females, height 171 ± 6 cm, body mass 65 ± 9 kg, age 27 ± 3 years) and one of older adults (G4, 5 males and 14 females, height 169 ± 8 cm, body mass 71 ± 12 kg, age 72 ± 6 years). All the participants completed a self-selected warm-up running on a treadmill, typically lasting between 3 and 5 min (*31*, *40*). After being instructed about the protocol, they completed a different set of measurements, depending on the protocol they were assigned to.

The experimental protocol E1 consisted of walking (at 1.40 m/s) and running (at 2.80 m/s) overground and on a treadmill. The speeds were chosen as the commonly reported average comfortable locomotion speeds (*29*, *40*). For the overground trials, we used a light-barrier system to control the speeds (average values of 1.40 ± 0.03 and 2.80 ± 0.04 m/s) in two consecutive sectors of 3 m each.

The experimental protocol E2 consisted of walking (1.10 m/s for females, 1.20 m/s for males) and running (2.00 m/s for females, 2.20 m/s for males) on one standard (Laufergotest, Erich Jäger, Würzburg, Germany) and one uneven-surface (Woodway^®^, Weil am Rhein, Germany, Fig. S1, Movie S1) treadmill (*7*). The uneven-surface treadmill’s belt consisted of terrasensa^®^ classic modules (Sensa^®^ by Huebner, Kassel, Germany). The speeds were chosen after a pilot study in which we estimated the average comfortable locomotion speed on the uneven-surface treadmill for males and females separately. Part of the data from this experimental protocol was previously reported (*7*).

The experimental protocol E3 consisted of walking (1.20 m/s for the group of young adults G3, 1.10 m/s for the group of old adults G4) on a treadmill (BalanceTutor™, MediTouch LTD, Netanya, Israel, Movie S1) that could provide mediolateral (through sudden displacement of the belt-supporting platform) and anteroposterior (through rapid acceleration of the belt) perturbations. The speeds were chosen after a pilot study in which we estimated the average comfortable walking speed under perturbed conditions for young and old adults separately. Both the perturbed and unperturbed trials lasted six minutes. The perturbed trials began with ~15 s of unperturbed locomotion. Afterwards, the participants were informed about the beginning of perturbations, which were delivered randomly (left or right mediolateral displacement or acceleration) every ~3 s (G3: 3.786 ± 0.986 s; G4: 3.072 ± 0.434 sec) at unspecified phases of the gait cycle. The interval between perturbations was a function of perturbation intensity (e.g. the larger the displacement of the platform, the longer the time needed to reset the controls and start a new perturbation). Perturbation intensities were set in the proprietary software on a scale from 1 to 30. Mediolateral perturbations were set at an intensity of 15 for G3 and 10 for G4, while the accelerations were set at 12 for G3 and 8 for G4. Intensities were chosen after a pilot study in which we defined the “mostly challenging intensity before failure” for young and old adults separately.

The protocols E2 and E3 both included external perturbations to locomotion. However, the timing (continuous in E2 and every 3 s in E3) and mechanics (uneven surface in E2 and displacement of the treadmill’s belt in E3) where of different nature. We chose two perturbation paradigms to allow for generalization of the outcomes. During the trials, the participants of both E2 and E3 were instructed to keep looking at a fixed spot in front of them and avoid looking at the treadmill’s belt.

### EMG recordings

Independently on the experimental protocol, the muscle activity of the following 13 ipsilateral (right side) muscles was recorded: gluteus medius (ME), gluteus maximus (MA), tensor fasciæ latæ (FL), rectus femoris (RF), vastus medialis (VM), vastus lateralis (VL), semitendinosus (ST), biceps femoris (long head, BF), tibialis anterior (TA), peroneus longus (PL), gastrocnemius medialis (GM), gastrocnemius lateralis (GL) and soleus (SO). The electrodes were positioned as extensively reported previously (*27*, *31*). After around 60 s habituation (*7*), we recorded one trial of 60 s for each participant with an acquisition frequency of 1 kHz (E2) or 2 kHz (E1 and E3) by means of a 16-channel wireless bipolar EMG system (E2: myon m320, myon AG, Schwarzenberg, Switzerland; E1 and E3: aktos, menios GmbH, Ratingen, Germany). For the EMG recordings, we used foam-hydrogel electrodes with snap connector (H124SG, Medtronic plc, Dublin, Ireland). The first 30 gait cycles of the recorded trial were considered for subsequent analysis (*31*). For the overground locomotion part of E1, due to limited length of the walkway (20 m), the participants were asked to repeat the trials 10 times for the subsequent concatenation of the data. Trials that did not match the target speed with a tolerance of ± 0.05 m/s in walking and ± 0.10 m/s in running were repeated.

### Gait cycle breakdown

The gait cycle breakdown was obtained by the elaboration of the data acquired by a 3D accelerometer operating at 148 Hz and synchronized with the EMG system. The accelerometer was strapped to the right shoe, over the most distal portion of the second to fourth metatarsal bones. The data was low-pass filtered using a 4^th^ order IIR Butterworth zero-phase filter with cut-off frequency of 15 Hz. For estimating touchdown, we used the modified foot contact algorithm developed by Maiwald et al. (*41*). For estimating lift-off, we adopted our foot acceleration and jerk algorithm (*7*). The jerk algorithm searches for the global maximum of the vertical acceleration between two consecutive touchdown events to estimate the lift-off (LOe, where the “e” stays for “estimated”). This estimation, however, does not provide an accurate identification of the lift-off and needs some refinement. To get closer to the “real” lift-off timing, a characteristic minimum in the vertical acceleration (i.e. when the jerk equals zero) of the foot is identified in a reasonably small neighborhood of the LOe. We found [LOe – 250 ms, LOe + 100 ms] for both walking and running to be the sufficiently narrow interval needed to make the initial lift-off estimation. However, we reduced this interval to [LOe – 150 ms, LOe + 100 ms] in the presence of external perturbations (i.e. for E2 and E3). Both the approaches for the determination of touchdown and lift-off have been validated using force plate data (AMTI BP600, Advanced Mechanical Technology, Inc., Watertown, MA, USA) from 15 participants walking and running overground at six different velocities without perturbations and from the data recorded in G1 overground. We then calculated the true errors between the contact times detected via force plate and those obtained from the acceleration data and used the averages to correct the calculations. Errors were of 1.9 ms for touchdown and 13.1 ms for lift-off in walking and −4.1 ms and −13.2 ms for running.

### Muscle synergies extraction

For the experimental protocol E1, the overground EMG recordings were concatenated after identification of the complete gait cycles (touchdown to touchdown of the right foot). For the other protocols, we used the 30 gait cycles per trial described above. Muscle synergies data were extracted through a custom script (R v3.5.3, R Found. for Stat. Comp.) using the classical Gaussian non-negative matrix factorization (NMF) algorithm (*7*, *29*, *31*, *42*). The raw EMG signals were band-pass filtered within the acquisition device (cut-off frequencies 10 and 500 Hz). Then the signals were high-pass filtered, full-wave rectified and lastly low-pass filtered using a 4^th^ order IIR Butterworth zero-phase filter with cut-off frequencies 50 Hz (high-pass) and 20 Hz (low-pass for creating the linear envelope of the signal) as previously described (*7*). After subtracting the minimum, the amplitude of the EMG recordings obtained from the single trials was normalized to the maximum activation recorded for every individual muscle (i.e. every EMG channel was normalized to its maximum in every trial) (*27*, *31*). Each gait cycle was then time-normalized to 200 points, assigning 100 points to the stance and 100 points to the swing phase (*7*, *27*, *30*, *31*). The reason for this choice is twofold (*31*). First, dividing the gait cycle into two macro-phases helps the reader understanding the temporal contribution of the different synergies, diversifying between stance and swing. Second, normalizing the duration of stance and swing to the same number of points for all participants (and for all the recorded gait cycles of each participant) makes the interpretation of the results independent from the absolute duration of the gait events. Synergies were then extracted through NMF as previously described (*7*, *31*). For the analysis, we considered the 13 muscles described above (ME, MA, FL, RF, VM, VL, ST, BF, TA, PL, GM, GL and SO). The m = 13 time-dependent muscle activity vectors were grouped in a matrix V with dimensions m × n (m rows and n columns). The dimension n represented the number of normalized time points (i.e. 200). The matrix V was factorized using NMF so that V ≈ V_R_ = WH. The new matrix V_R_, reconstructed multiplying the two matrices W and H, approximates the original matrix V. The motor primitives (*29*, *43*) matrix H contained the time-dependent coefficients of the factorization with dimensions r × n, where the number of rows r represents the minimum number of synergies necessary to satisfactorily reconstruct the original set of signals V. The motor modules (*29*, *44*) matrix W, with dimensions m × r, contained the time-invariant muscle weightings, which describe the relative contribution of single muscles within a specific synergy (a weight was assigned to each muscle for every synergy). H and W described the synergies necessary to accomplish the required task (i.e. walking or swimming). The update rules for W and H are presented in Equation (S1) and Equation (S2).

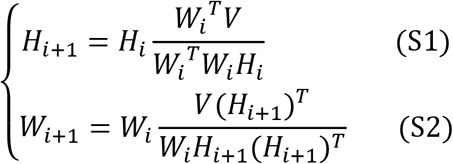

The quality of reconstruction was assessed by measuring the coefficient of determination R^2^ between the original and the reconstructed data (V and V_R_, respectively). The limit of convergence for each synergy was reached when a change in the calculated R^2^ was smaller than the 0.01% in the last 20 iterations (*29*) meaning that, with that amount of synergies, the signal could not be reconstructed any better. This operation was first completed by setting the number of synergies to 1. Then, it was repeated by increasing the number of synergies each time, until a maximum of 10 synergies. The number 10 was chosen to be lower than the number of muscles, since extracting a number of synergies equal to the number of measured EMG activities would not reduce the dimensionality of the data. Specifically, 10 is the rounded 75% of 13, which is the number of considered muscles. For each synergy, the factorization was repeated 10 times, each time creating new randomized initial matrices W and H, in order to avoid local minima (*45*). The solution with the highest R^2^ was then selected for each of the 10 synergies. To choose the minimum number of synergies required to represent the original signals, the curve of R^2^ values versus synergies was fitted using a simple linear regression model, using all 10 synergies. The mean squared error (*46*) between the curve and the linear interpolation was then calculated. Afterwards, the first point in the R^2^-vs.-synergies curve was removed and the error between this new curve and its new linear interpolation was calculated. The operation was repeated until only two points were left on the curve or until the mean squared error fell below 10^-4^. This was done to search for the most linear part of the R^2^-versus-synergies curve, assuming that in this section the reconstruction quality could not increase considerably when adding more synergies to the model.

### Local dynamic stability of motor primitives

We assessed the local dynamic stability of motor primitives using the maximum finite-time Lyapunov exponents (MLE) (*19*). Usually MLE are extracted after reconstruction of the state space through delay-coordinate embedding starting from a measured one-dimensional time series (*35*). The state space is a set of all the possible states of a system at any given time, the variables of which might be position, velocity, temperature, color, species, voltage and many others (*1*, *5*, *35*). Yet, in our typical experimental setups involving complex living systems like humans, the state space is often unknown. Theoretically, the behavior of a purely chaotic dynamical system can be predicted by using only a small set of observations on its state (e.g. joint angles, or kinematics, or accelerations, etc.) without losing information on its properties (*7*, *20*). For this reason, it is common to use the recorded data to reconstruct the state space, usually by means of the “delay embedding theorem” (*47*). Typically MLE are then calculated from data expanded in the state space (*5*, *35*). We avoided this passage by assuming that the space we are interested into had dimension equal to the factorization rank (i.e. the minimum number of synergies necessary to sufficiently reconstruct the original EMG signals). The work from Sauer and colleagues (*48*) allows to use n-dimensional measurements instead of the classical state space reconstruction. For instance, if one trial was factorized by NMF into four synergies, we would calculate the MLE of the resulting four motor primitives using four as embedding dimension. This approach has the advantage that it does not require the estimation of a suitable delay and embedding dimension, the latter being particularly sensitive to the presence of noise, which is very likely in experimental data (*49*). The motor primitives associated to the fundamental synergies extracted using the methods described above, were analyzed as follows. Each set of motor primitives was a time series of 30 gait cycles, normalized in time as described above (100 points for the stance and 100 for the swing, for a total of 6000 points per trial). Primitives were then scaled to have the same variance by subtracting the mean and dividing by the standard deviation in order to avoid having different dynamical ranges across the data set (*50*). For every point x_i_ in each time series (or set of motor primitives), we searched for the nearest neighbors of the point x_i_ excluding the neighborhood points [x_i − 100_, x_i + 100_]. This interval was chosen in order to impose a temporal separation between the nearest neighbors (Theiler window), making sure that they were on different trajectories (or gait cycles), as previously described (*19*). Once the algorithm found the nearest neighbors, we proceeded to calculate the logarithm of the divergence between the trajectory of each point and its nearest neighbor’s and for a maximum of 300 consecutive time points. For each trial, the divergence curve was calculated as the average of all divergence curves obtained from each point in the time series and their neighbors (*19*). We then defined MLE as the slope of the most linear part of the divergence curve, starting from the first point. To define linearity, we imposed the R^2^ between the curve and its linear interpolation to be bigger than 0.9. Across all trials for each experimental setup, we then found the minimum number of points needed to reach a linear interpolation with R^2^ > 0.9 and used these values to recalculate the final MLE. The minimum number of points was three in E1, E2 and E3 and this is the value we used; the maximum was 8 (E1), 9 (E2), and 7 (E3), with average values of 5.3 ± 1.1 (E1), 5.5 ± 1.1 (E2), 5.2 ± 0.8 (E3).

### Width of motor primitives

We compared motor primitives by evaluating the full-width at half maximum (FWHM), a metric useful to describe the duration of activation patterns (*7*, *24*, *27*). The FWHM was calculated cycle-by-cycle as the number of points exceeding each cycle’s half maximum, after subtracting the cycle’s minimum and then averaged (*24*). The FWHM was calculated only for the motor primitives relative to fundamental synergies. A fundamental synergy can be defined as an activation pattern whose motor primitive shows a single main peak of activation (*7*). When two or more fundamental synergies are blended into one, a combined synergy appears. Combined synergies usually constitute, in our data, 10 to 20% of the total extracted synergies. Due to the lack of consent in the literature on how to interpret them, we excluded the combined synergies from the FWHM analysis. The recognition of fundamental synergies was carried out based on a previously reported approach (*29*, *30*), which involves the creation of a training set and the subsequent supervised clustering of similar primitives.

### Statistics

To investigate the main effects on the MLE and FWHM of locomotion type (i.e. walking or running), condition (i.e. overground or treadmill), perturbations and age, we fitted the data using a generalized linear model with Gaussian error distribution. The homogeneity of variances was tested using the Levene’s test. If the residuals were normally distributed, we carried out a two-way repeated measures ANOVA with type II sum of squares, the independent variables being: locomotion type (walking or running) and condition (overground or treadmill) in E1; locomotion type (walking or running) and condition (perturbed or unperturbed) in E2; locomotion condition (perturbed or unperturbed) and age (young or old) in E3. If the normality assumptions on the residuals were not met, we used a robust (rank-based) ANOVA from the R package Rfit (function “raov”) (*51*, *52*). All the significance levels were set to α = 0.05 and the statistical analyses were conducted using R v3.5.3 (R Found. for Stat. Comp.).

### Data availability

In the supplementary Data S1, available upon request, we made available: a) the metadata with anonymized participant information, b) the raw EMG, c) the touchdown and lift-off timings of the recorded limb, d) the filtered and time-normalized EMG, e) the muscle synergies extracted via NMF and f) the code to process the data, including the scripts to calculate the MLE of motor primitives.

The file “participants_data.dat” is available in ASCII and RData (R Found. for Stat. Comp.) format and contains:

- Code: the participant’s code
- Experiment: the experimental setup in which the participant was involved (E1 = walking and running, overground and treadmill; E2 = walking and running, even- and uneven-surface; E3 = unperturbed and perturbed walking, young and old)
- Group: the group to which the participant was assigned (see above for the details)
- Sex: the participant’s sex (M or F)
- Speed: the speed at which the recordings were conducted in [m/s]
- Age: the participant’s age in years (participants were considered old if older than 65 years, but younger than 80)
- Height: the participant’s height in [cm]
- Mass: the participant’s body mass in [kg].

The files containing the gait cycle breakdown are available in ASCII and RData (R Found. for Stat. Comp.) format. The files are structured as data frames with 30 rows (one for each gait cycle) and two columns. The first column contains the touchdown incremental times in seconds. The second column contains the duration of each stance phase in seconds. Each trial is saved both as a single ASCII file and as an element of a single R list. Trials are named like “CYCLE_TIMES_P0020,” where the characters “CYCLE_TIMES” indicate that the trial contains the gait cycle breakdown times and the characters “P0020” indicate the participant number (in this example the 20th).

The files containing the raw, filtered and the normalized EMG data are available in ASCII and RData (R Found. for Stat. Comp.) format. The raw EMG files are structured as data frames with 30000 rows (one for each recorded data point) and 14 columns. The first column contains the incremental time in seconds. The remaining thirteen columns contain the raw EMG data, named with muscle abbreviations that follow those reported in the Materials and Methods section of this Supplementary Materials file. Each trial is saved both as a single ASCII file and as an element of a single R list. Trials are named like “RAW_EMG_P0053”, where the characters “RAW_EMG” indicate that the trial contains raw emg data and the characters “P0053” indicate the participant number (in this example the 53rd). The filtered and time-normalized emg data is named, following the same rules, like “FILT_EMG_P0053”.

The files containing the muscle synergies extracted from the filtered and normalized EMG data are available in ASCII and RData (R Found. for Stat. Comp.) format. The muscle synergies files are divided in motor primitives and motor modules and are presented as direct output of the factorization and not in any functional order. Motor primitives are data frames with a number of rows equal to the number of synergies (which might differ from trial to trial) and 6000 columns. The rows contain the time-dependent coefficients (motor primitives), one row for each synergy (named e.g. “Syn1, Syn2, Syn3”, where “Syn” is the abbreviation for “synergy”). Each gait cycle contains 200 data points, 100 for the stance and 100 for the swing phase which, multiplied by the 30 recorded cycles, result in 6000 data points distributed in as many columns. Each set of motor primitives is saved both as a single ASCII file and as an element of a single R list. Trials are named like “SYNS_H_P0012”, where the characters “SYNS_H” indicate that the trial contains motor primitive data and the characters “P0012” indicate the participant number (in this example the 12th). Motor modules are data frames with 13 rows (number of recorded muscles) and a number of columns equal to the number of synergies (which might differ from trial to trial). The rows, named with muscle abbreviations that follow those reported in the Materials and Methods section of this Supplementary Materials file, contain the time-independent coefficients (motor modules), one for each synergy and for each muscle. Each set of motor modules relative to one synergy is saved both as a single ASCII file and as an element of a single R list. Trials are named like “SYNS_W_P0082”, where the characters “SYNS_W” indicate that the trial contains motor module data and the characters “P0082” indicate the participant number (in this example the 82nd).

All the code used for the preprocessing of EMG data, the extraction of muscle synergies and the calculation of MLE is available in R (R Found. for Stat. Comp.) format. Explanatory comments are profusely present throughout the scripts (“SYNS.R”, which is the script to extract synergies, “fun_synsNMFn.R”, which contains the NMF function, “MLE.R”, which is the script to calculate MLE of motor primitives and “fun_MLE.R”, which contains the MLE function).

---

**Fig. S1.**
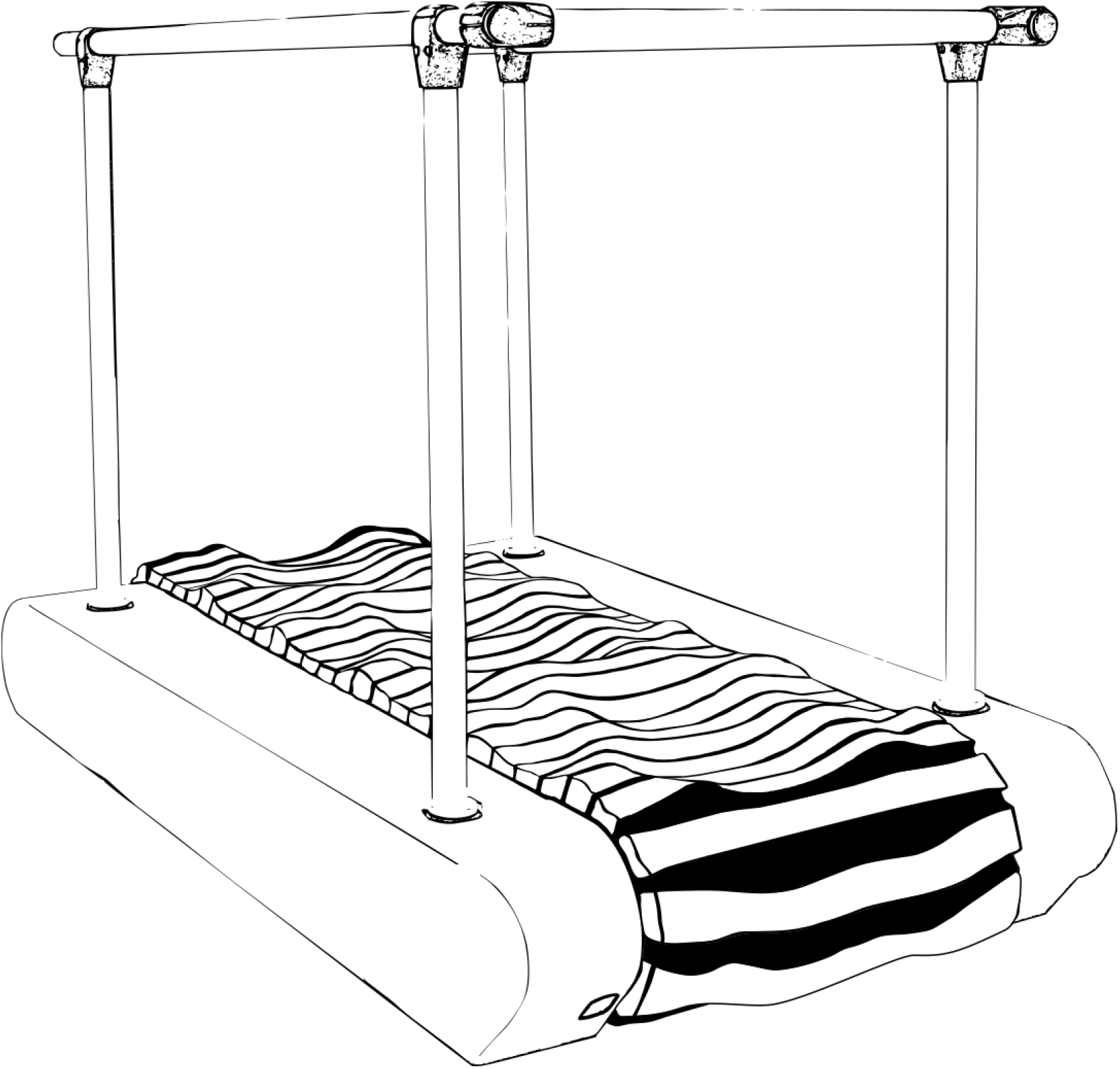
Sketch of the uneven-surface treadmill used for the experimental protocol E2. The belt of this treadmill was built to reproduce an uneven-terrain environment.

**Fig. S2.**
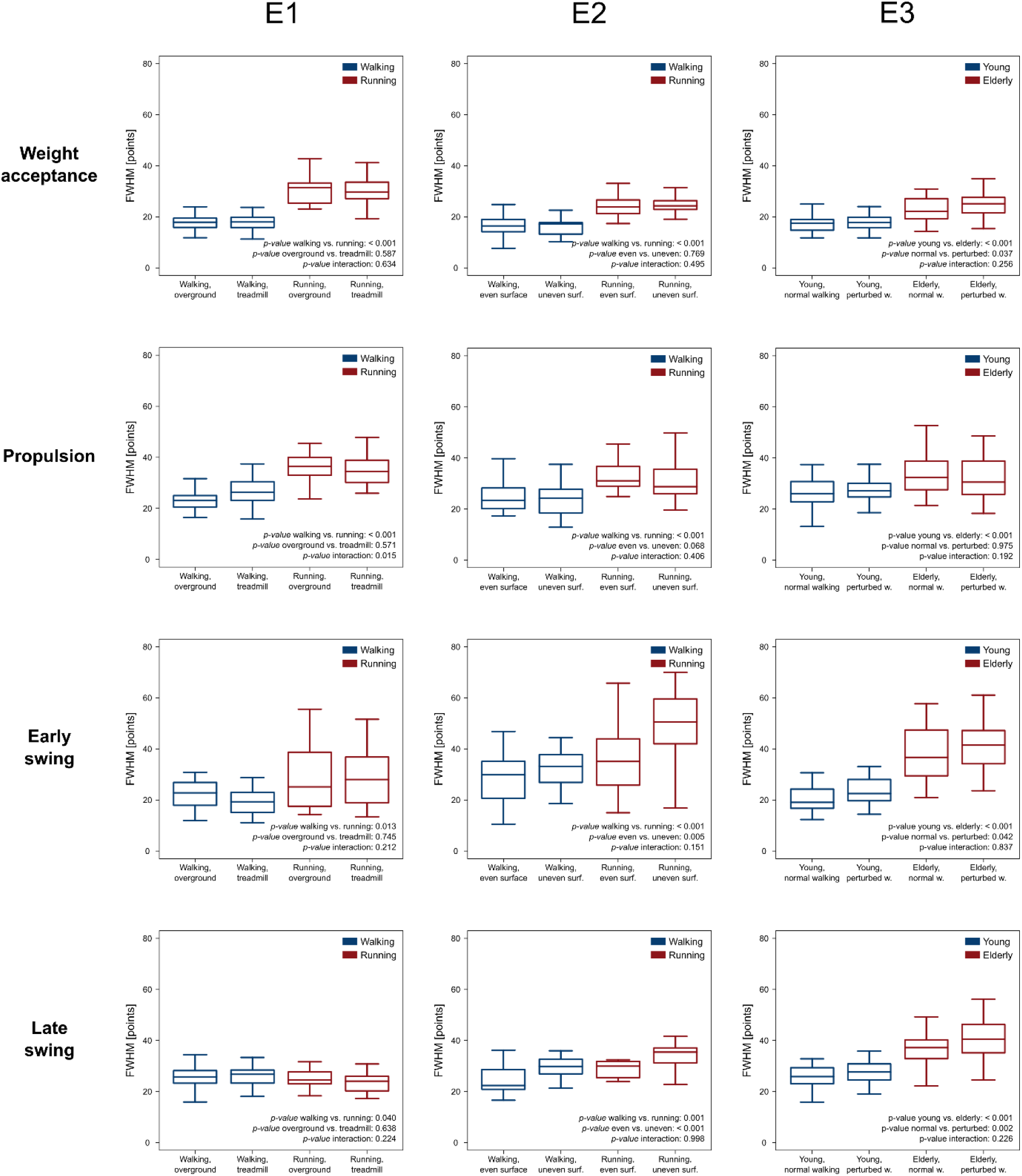
Full width at half maximum of motor primitives not shown in Table 1. Boxplots describing the full width at half maximum (FWHM) of the motor primitives extracted from the data of the three experimental setups (E1 = walking and running, overground and treadmill; E2 = walking and running, even- and uneven-surface; E3 = unperturbed and perturbed walking, young and old). Motor primitives are the temporal coefficients of the four fundamental synergies for locomotion. Lower FWHM imply shorter duration of activation.

**Fig. S3.**
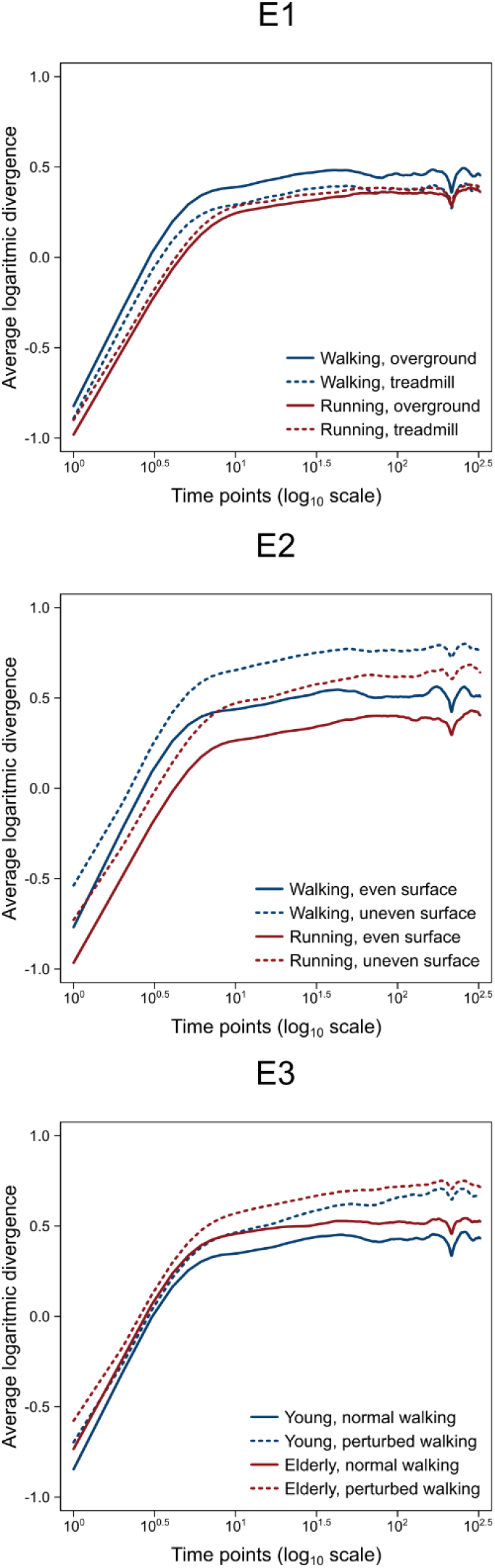
Average logaritmic divergence curves of motor primitives with their original vertical intercept not shown in Fig. 3. Curves describing the average logarithmic divergence curves for the three experimental setups (E1 = walking and running, overground and treadmill; E2 = walking and running, even- and uneven-surface; E3 = unperturbed and perturbed walking, young and old).

**Movie S1. Treadmills for perturbed locomotion used in this study.** This video shows the typical setup of the two treadmills used for introducing external perturbations during locomotion. The first treadmill is equipped with an uneven-surface belt. The second one can provide sudden accelerations of the belt and displacements of the platform to induce anteroposterior and mediolateral perturbations, respectively.

**Data S1. (separate file) Raw electromyographic (EMG) data and code to extract muscle synergies and calculate Maximum Lyapunov Exponents (MLE).** This supplementary data set contains: a) the metadata with anonymized participant information, b) the raw EMG acquired during locomotion, c) the touchdown and lift-off timings of the recorded limb, d) the filtered and time-normalized EMG, e) the muscle synergies extracted via NMF and f) the code written in R (R Found. for Stat. Comp.) to process the data, including the scripts to calculate the MLE of motor primitives.

## Acknowledgments

We thank Juri Taborri for the tireless contribution to different parts of the measurements and are grateful to all the participants that showed great commitment and interest during the experiments. We disclose any professional relationship with companies or manufacturers who might benefit from the results of the present study;

## Author contributions

Conceptualization: A.Sa., L.B., A.E., M.S., and A.A.; Data curation: A.Sa., A.E., N.E.; Formal analysis: A.Sa.; Investigation: A.Sa., L.B., A.E. and N.E.; Methodology: A.Sa., A.E., A.Sc., and A.A.; Project administration: A.Sa., A.K., M.S., and A.A.; Resources: A.Sa., L.B., N.E., A.K., M.S., and A.A.; Software: A.Sa.; Supervision: A.Sa., and A.A.; Validation: A.Sa.; Visualization: A.Sa.; Writing – original draft: A.Sa., and A.A.; Writing – review & editing: A.Sa., L.B., A.E., A.Sc., N.E., A.K., M.S., and A.A.;

## Data and materials availability

All data is available in the main text or the supplementary materials.

